# Abscisic acid inhibits germination of Striga seeds and is released by them as a rhizospheric signal providing competitive advantage and supporting host infestation

**DOI:** 10.1101/2023.07.06.548005

**Authors:** Muhammad Jamil, Yagiz Alagoz, Jian You Wang, Guan-Ting Erica Chen, Lamis Berqdar, Najeh M. Kharbatia, Juan C. Moreno, Hendrik N. J. Kuijer, Salim Al-Babili

**Affiliations:** The BioActives Lab., Center for Desert Agriculture, King Abdullah University of Science and Technology, Thuwal 23955-6900, Saudi Arabia; Plant Science Program, Biological and Environmental Science and Engineering (BESE) Division, King Abdullah University of Science and Technology (KAUST), Thuwal 23955-6900, Saudi Arabia; Analytical Chemistry Core Lab, King Abdullah University of Science and Technology (KAUST), Thuwal 23955-6900, Saudi Arabia

**Keywords:** Abscisic acid, Conditioning, Dormancy, Fluridone, Striga, Strigolactones

## Abstract

Seeds of the root parasitic plant *Striga hermonthica* undergo a conditioning process under humid and warm environments before germinating in response to host-released stimulants, particularly strigolactones (SLs). The plant hormone abscisic acid (ABA) regulates different growth and developmental processes, and stress response; however, its role during Striga seed germination and early interactions with host plants is under-investigated.Here, we show that ABA inhibited Striga seed germination and that hindering its biosynthesis induced conditioning and germination in unconditioned seeds, which was significantly enhanced by treatment with the SL analog *rac*-GR24. However, the inhibitory effect of ABA remarkably decreased during conditioning, confirming the loss of sensitivity towards ABA in later developmental stages. ABA measurement showed a significant reduction of its content during the early conditioning stage and a significant increase upon *rac*-GR24-triggered germination. We observed this increase also in released seed exudates, which was further confirmed by using the Arabidopsis ABA-reporter GUS marker line.Seed exudates of germinated seeds, containing elevated levels of ABA, impaired the germination of surrounding Striga seeds *in vitro* and promoted root growth of a rice host towards germinated Striga seeds. Application of ABA as a positive control caused similar effects, indicating its function in Striga/Striga and Striga/host communications.In summary, we show that ABA is an essential player during seed dormancy and germination processes in Striga and acts as a rhizospheric signal released by germinated parasitic seeds to provide a competitive advantage and support host infestation.

**Societal Impact Statement:** The root parasitic plant *Striga hermonthica* is a severe threat to cereal’s yield, endangering global food security. Herein, we uncover a new role of the known plant hormone abscisic acid (ABA) as a rhizospheric signal released by germinated Striga seeds, allowing them to better compete with surrounding un-conditioned seeds and facilitating host infestation. Our findings can help in developing strategies to control this parasite and mitigate its negative impact on the food supply and income of smallholder farmers.

## 1 INTRODUCTION

The root parasitic plant *Striga hermonthica* (purple witchweed) is one of the major biological threats to global food security, impacting cereal production, particularly in Africa (Spallek et al., 2013; Westwood, 2013). Several factors, including subterranean damage, seed longevity and dormancy, complex life cycle, and the dependency on host-released germination stimulants, have made Striga control a very challenging task (Samejima et al., 2016; Kountche et al., 2019; Jamil et al., 2021). Striga seeds can remain viable for up to 15 years in soil, waiting for favourable germination conditions, including the presence of a suitable host (Ejeta and Gressel, 2007). Indeed, Striga seed germination occurs only when suitable host is present in close vicinity, which is ensured by the requirement for host-released stimulants, mainly strigolactones (SLs) (Yoneyama et al., 2010). Most parasitic seeds, including those of *Orobanche, Phelipanche*, and *Striga* species, need to be exposed to special enviroments for a period of time to become able to perceive host-released signals that trigger their germination (Matusova et al., 2004). This stage, generally called conditioning, has been reported to be associated with global genome demethylation in the root parasitic plant *Phelipanche ramosa* (Lechat et al., 2015). In the case of Striga seeds, the conditioning requires temperature of around 30°C and moist environment for one to two weeks (Matusowa et al., 2004). Perception of host released SLs by conditioned seeds leads to their germination, which is followed by developing a radicle that differentiates into a specialized organ, the ‘haustorium’, to siphon off water and nutrients from the host, which promotes the growth of the parasite that emerges above the surface, flowers and produces a large number of seeds building up a persistent seed bank in infested areas (Hearne, 2009). Despite substantial progress in understanding Striga seed germination, our knowledge about the internal programming, particularly with respect to hormonal balance in Striga seeds during pre-conditioning and germination process, remains vague.

Abscisic acid (ABA) is a major determinant of seed dormancy and an inhibitor of seed germination in many plant species (Kucera et al., 2005; Shu et al., 2016). One of the possibilities to determine the role of ABA in Striga parasitism is to suppress ABA biosynthesis in the seeds. Application of carotenoid biosynthesis inhibitors such as 1-methyl-3-phenyl-5-[3-trifluoromethy1-(phenyl)]-4-(lH)-pyridinone (fluridone) and norflurazon, have been widely used to inhibit ABA biosynthesis by reducing the availability of its carotenoid-precursor. FL, an inhibitor of phytoene desaturase that converts phytoene to phytofluene in the upstream carotenoid biosynthesis pathway (Lee et al., 2015), ultimately inhibits ABA biosynthesis (Yamagishi et al., 2009). Moreover, the carotenoid biosynthesis inhibitor fluridone (FL) regulates seed dormancy and germination, presumably by decreasing ABA formation in *Orobanche minor* and *Striga asiatica* seeds (Chae et al., 2004; Kusumoto et al., 2006). FL application shortens the conditioning period and enhances the responsiveness to germination stimulants (Chae et al., 2004; Jamil et al., 2023). These results indicate a role of ABA in Striga seed conditioning and germination and point to the presence of functional ABA biosynthesis and perception. It can be assumed that fluridone-induced seed germination occurs via SL-independent signaling pathway (Machin and Bennett, 2020).

In this work, we demonstrate that Striga seed dormancy and germination are closely associated with endogenous ABA levels. We show that Striga seeds released ABA upon germination, which impaired the germination of neighboring Striga seeds. ABA released by germinated Striga seeds also attracted the growth of host roots towards germinated Striga. Our results reveal an overlooked function of ABA as a rhizospheric signal in communication within the Striga seed population and between arising Striga seedlings and host plants. Moreover, they open up the possibility of using inhibitors of ABA biosynthesis as a tool to control Striga infestation by inducing lethal seed germination in the host’s absence, i.e. suicidal germination (Zwanenburg et al., 2016; Brun et al., 2018).

## 2 MATERIALS AND METHODS

### 2.1 Hormonal profiling

For hormone quantification, we followed the protocol published by Wang et al., (2022). In brief, we spiked around 20 mg ground Striga seeds with internal standards D_6_-ABA (1.0 ng) in 1.5 ml methanol, sonicated the mixture for 15 min in an ultrasonic bath (Branson 3510 ultrasonic bath), then centrifuged it for 10 min at 4,000 x g at 4°C. The supernatant was collected, and the pellet was re-extracted with 1.5 ml of the same solvent. Then, the two supernatants were combined and dried under a vacuum. The sample was re-dissolved in 150 μl of acetonitrile:water mixture (25:75, v:v) and filtered through a 0.22 μm filter for LC-MS analysis. To quantify plant hormones in the seed exudate, 200 μl collected exudate was spiked with internal standards D_6_-ABA (1.0 ng) and filtered through a 0.22 μm filter for LC-MS analysis. Plant hormones were analyzed using an HPLC-Q-Trap-MS/MS with Multiple Reaction Monitoring (MRM) mode. Chromatographic separation was achieved on a ZORBAX Eclipse plus C_18_ column (150 × 2.1 mm; 3.5 μm; Agilent). Mobile phases consisted of water:acetonitrile (95:5, v:v) and acetonitrile, both containing 0.1% formic acid. A linear gradient was optimized as follows (flow rate, 0.4 ml/min): 0−17 min, 10% to 100% B, followed by washing with 100% B and equilibration with 10% B. The injection volume was 10 μl, and the column temperature was maintained at 40 C for each run. Mass spectrometry was conducted in electrospray and MRM mode in negative ion mode. Relevant instrumental parameters were set as follows: ion source of turbo spray, ion spray voltage of (-) 4500 V, curtain gas of 25 psi, collision gas of medium, gas 1 of 45 psi, gas 2 of 30 psi, turbo gas temperature of 500 C, entrance potential of -10 V. The characteristic MRM transitions (precursor ion → product ion) were 263.2→153.1 for ABA; 269.2→159.1 for D_6_-ABA.

### 2.2 Striga seed germination bioassays

The Striga seeds used in this study were collected from sorghum infested fields near Wad Medani, Sudan in 2020. The germination bioassays to examine the effect of FL and or/ABA ± *rac*-GR24 (see Figure 1a) on the unconditioned Striga seeds were performed according to published protocol (Wang et al., 2022). After surface sterilization, the Striga seeds were dried and spread (50-100) uniformly on 9 mm glass fiber filter paper discs. Then two filter papers were added to a 9 cm Petri plate, and five unconditioned Striga seed discs were transferred to each plate. Each plate was treated with 4 ml FL alone (100 μM), *rac*-GR24±FL (1.0 μM±100 μM), or *rac*-GR24+FL+ABA (1.0 μM+100+100 μM). The plates were sealed with parafilm and incubated, and scanned from day 1 to day 7. After scanning and counting germinated and non-germinated seeds with SeedQuant (Braguy et al., 2021), the percentage germination was calculated.

**FIGURE 1.**
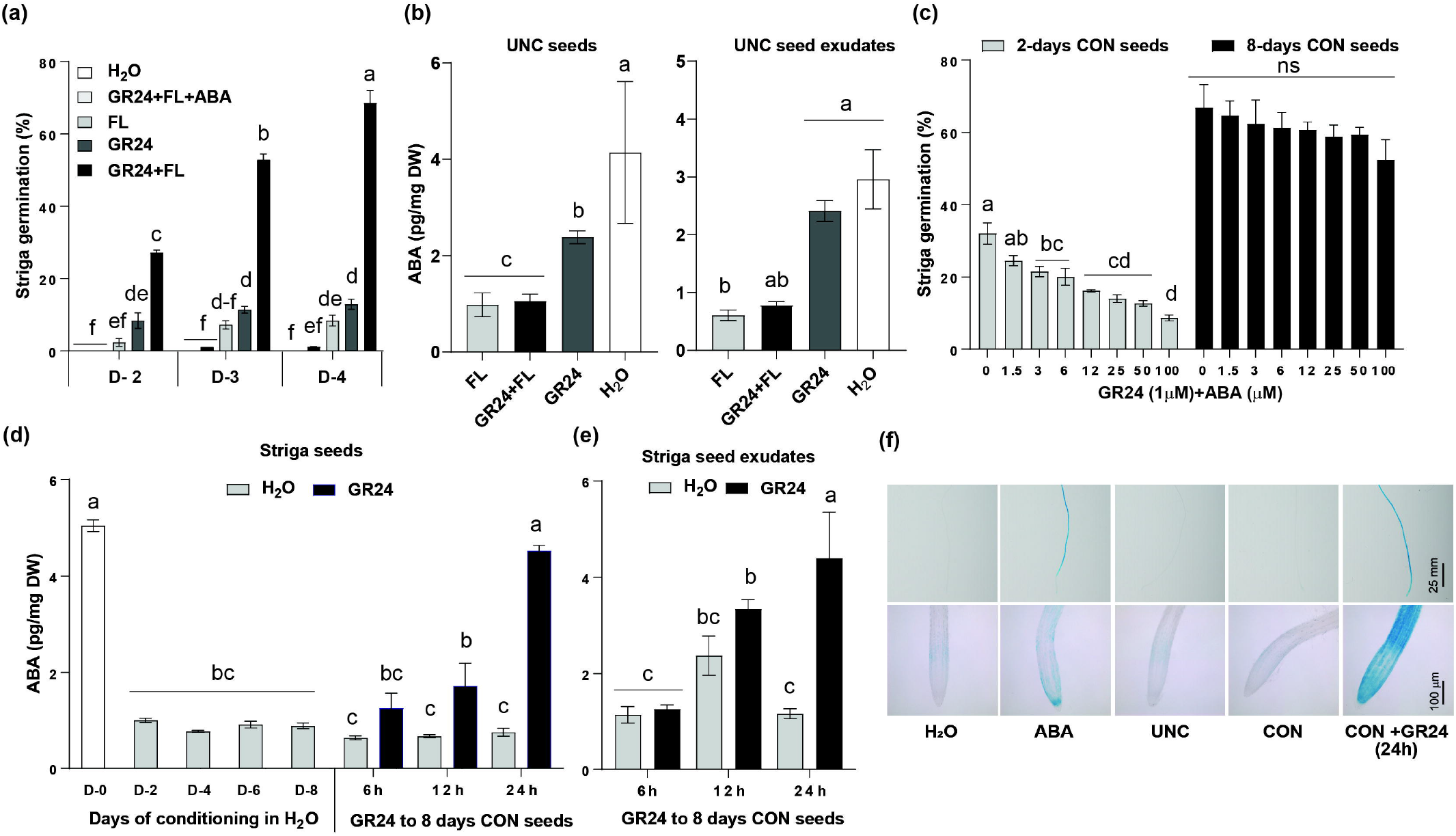
ABA role and content during conditioning and germination of Striga seeds. (a) Germination of unconditioned Striga seeds in response to Fluridone (FL) ±*rac*-GR24 treatment. FL (100 μM), *rac-*GR24 (1.0 μM) and ABA (100 μM) were applied on unconditioned Striga seeds (3 ml per plate) on day 0, and Striga seeds were scanned from day 1 to day 4 to determine their germination rate. Data are means ±SE (n=5). (b) ABA content in unconditioned Striga seeds and seed exudates in response to FL ±*rac*-GR24 treatment. Striga seeds were treated with FL (100 μM) ±*rac*-GR24 (1.0 μM) for 24 h, and ABA levels were measured in unconditioned Striga seeds and seed exudates. Data are means ±SE (n=3). (c) Effect of ABA on Striga seed germination during the conditioning period. Striga seeds were conditioned in water for two or eight days. *rac*-GR24 (1.0 μM) ±ABA (1.5, 3, 6, 12, 25, 50, and 100 μM) was applied for 24 h, and germinated and non-germinated seeds were counted. Data are means ±SE (n=6). (d-e) Kinetics of ABA content in Striga seeds and seed exudates during conditioning and germination processes. Conditioned Striga seeds were treated with *rac*-GR24 (1.0 μM), and ABA was quantified by LC-MS in Striga seeds and seed exudates. Data are means ±SE (n=3). (f) GUS staining analysis of an Arabidopsis ABA reporter line upon exposure to ABA released by germinated Striga seeds. Exudates of unconditioned, conditioned and *rac*-GR24-treated (1.0 μM, 24h) and conditioned seeds were applied to roots of the Arabidopsis *pMKKK18::GUS* ABA marker line for six hours, followed by GUS-staining. We used water and ABA (50 μM) as a negative and positive control, respectively. We detected a strong GUS signal upon the application of exudates from *rac*-GR24-treated seeds and of ABA. Data are means ±SE (n=4). The annotation with various letters denotes significance (one-way ANOVA, Tukey’s post hoc test, P <0.05). ABA: Abscisic acid; UNC: unconditioned; CON: Conditioned; FL: Fluridone.

In another experiment (Figures 3a and 3b), Striga seeds were surface sterilized by 2% sodium hypochlorite (NaClO) for seven minutes, followed by six subsequent washings with sterilized Mili Q water under a laminar flow cabinet. The Striga seeds were air-dried and spread (50-100) uniformly on 9 mm glass fiber filter paper discs. Then 12 discs with Striga seeds were transferred in a 9 cm Petri plates on a Whatman filter paper, moistened with 3 ml sterilized Mili Q water. The Petri plates were sealed with parafilm and covered with aluminum foil. The Striga seeds were incubated at 30 C for eight days for the conditioning process. These eight days conditioned seeds were treated with H_2_O and *rac*-GR24 (1.0 μM) to induce germination for 24 h to be used as the inducer discs. The next day, four discs containing new Striga seeds (pre-conditioned for 0 to 7 days) were put in the middle of a petri plate as assay discs. One inducer disc containing non-germinated seeds, germinated seeds or ABA alone were added in between these four discs. Then next day, the Striga inducer disc was removed, and all the assay discs were further treated with *rac*-GR24 (1.0 μM) and incubated for another 24 h to observe germination. This experiment was repeated using assay discs with seeds conditioned for 0 to seven days.

### 2.3 Striga emergence in pots under greenhouse conditions

A pot experiment using suicidal germination approach (Jamil et al., 2018) was conducted to determine the impact of SL ±FL. For this purpose, a mixture of soil (Stender), sand and rock at 3:1:1 ratio was prepared. About 0.5 L blank soil was added to the bottom of each 3 L plastic pots. Then about 20 mg (∼6000) Striga seeds were mixed thoroughly with 1.5 L soil mixture and added on top of blank soil in the same plastic pots. The pots were treated for two times (day 1 and day 3) with a previously described (Jamil et al., 2018; Jamil et al., 2022b), formulated SL analogs MP3 27EC ±FL (1.0+1.0 μM) or the corresponding blank solution. About three weeks after treatment, one-week-old susceptible rice (cv IAC 165) seedlings (four per pot) were planted in each pot. The number of emerged Striga seedlings was counted in each pot after 10 weeks of rice planting.

### 2.4 Host root/shoot growth experiments in rhizotron

Rice seeds (*Oryza sativa* L. cv IAC-165) were surface sterilized with 50% sodium hypochlorite (Milli-Q water to commercial bleach; 1:1 v/v) for 10 minutes, followed by 10 subsequent washings with sterilized MiliQ water in the laminar flow cabinet. The rice seeds were then transferred to magenta boxes filled with ½ MS + 4% (g/L) agar. Thereafter, the magenta boxes were kept in the dark in a 30°C incubator for 24 h. We then transferred all the magenta boxes to a light growth chamber with a day/night temperature of 28/22 C for seven days. The germinated seedlings were separated into two groups (with or without pre-germinated Striga seeds), and each seedling was transferred to a 9.5 cm square Petri dishes filled with agar under three different conditions: mock (50 ml of half MS + 7% agar), ABA (50 mL of 1/2 MS + 7% agar with 5 μM ABA), and mock/ABA, where the main root should be placed perpendicularly straight to the bottom of the dish. In mock/ABA plates, rice plants were placed on the mock side. After planting the seedlings, pre-germinated Striga seeds were spread on the upper right corner of the dish. To prevent ABA degradation, the dishes were covered with aluminum foil and placed in the light growth chamber for seven days. Root length, shoot length, crown root numbers, and lateral root numbers of the primary root toward the mock or ABA sides were recorded.

### 2.5 GUS staining bioassays and imaging

Seeds of the Arabidopsis *pMKKK18::GUS* marker line were surface sterilized with 20% household bleach and stratified for the consecutive 3 days before sowing on 1/2 MS agar plates. Seedlings were grown in a controlled growth cabinet at 21 C with 60% humidity under a long-day (16 h light/8 h dark) photoperiod. Five days old Arabidopsis seedlings were dipped in 24 wells plates containing 0.5 ml each of H_2_O, ABA (50 μM), and Striga seed exudates collected from unconditioned or conditioned Striga seeds ±*rac*-GR24 (1.0 μM). The exudates were prepared directly by adding 0.5 ml H_2_O on top of Striga seed discs (each containing ∼100 seeds) to dissolve the exudate released on paper discs. After 6 h of incubation at room temperature, seedlings were transferred into another 24-well plate containing GUS staining buffer. The GUS staining assays were performed according to the published work (Ablazov et al., 2022). After staining, images were collected via a professional DSLR and/or microscope.

### 2.6 Gene expression analysis

The total RNA was extracted from the Striga seeds using the Direct-zol™ RNA Miniprep Plus kit (Zymo Research) according to the manufacturer’s instructions. A 1.0 μg of total RNA was used for the cDNA synthesis using the iScript cDNA Synthesis kit (Bio-Rad). The qRT-PCR analysis was performed using CFX 384 Touch RT-PCR detection system with a reaction mixture of 2.0 μl of primer mix (2.0 μM from each F & R primer), 1.0 μl 1/5 diluted cDNA template, 5.0 μl SsoAdvanced™ Universal SYBR® Green Supermix (Bio-Rad), and 2.0 μl of dH2O to make a total volume of 10 μl. Three technical replicates for each of the four biological replicates were used to perform the experiments for each control and treatment condition. The relative quantification of gene expression was calculated according to the 2^−ΔΔct^ method (Livak et al., 2001). *ShTUB5* and *ShPP2A* were used as housekeeper reference controls for the relative gene expression analysis (Fernández-Aparicio et al., 2013). All primers used in this study are listed in **Table S1**. Statistical analyses were performed using Two-Way ANOVA (Tukey’s multiple comparison test).

## 3 RESULTS & DISCUSSION

### 3.1 ABA inhibits seed germination and delays conditioning of Striga seeds

Conditioning is the stratification of parasitic plant seeds under moist and warm environment for a specific period of time, which is required for them to become responsive to the germination stimulants released by host plants (Matusova et al., 2004); however, the conditioning time largely depends on the parasite species (Brun et al., 2018). To examine the function of ABA in Striga seed dormancy and conditioning, we first applied FL in the absence or presence of the SL analog *rac*-GR24 to unconditioned Striga seeds and determined their germination ratio on the 2^nd^, 3^rd^, and 4^th^ day after application (DAA). Treatment with FL (100 μM) or *rac*-GR24 (1.0 μM) alone caused 8-13% germination of unconditioned seeds on the 4 DAA. The combined application of FL + *rac*-GR24 (100+1.0 μM) boosted the germination of unconditioned seeds to around 70% on the 4^th^ DAA (Figure 1a; Figure S1), suggesting a synergetic effect of inhibiting ABA biosynthesis and triggering SL response. As expected, adding ABA (100 μM) to FL + *rac*-GR24 significantly decreased the germination rate to around 1.0% (Figure 1a). In general, we observed an increase in the germination rate over time and detected the highest values on the 4^th^ DAA. These data indicate that a proportion of the tested seeds have become conditioned even within one day and that inhibiting ABA biosynthesis alone is sufficient to trigger seed germination. To initiate germination, Striga seeds generally perceive SLs by HYPOSENSITIVE to LIGHT (ShHTL) receptors, the diverged family of α/β hydrolase-fold proteins in Striga (Tsuchiya & McCourt, 2015). After binding and acting via ShHTLs, the hydrolysis of SLs leads to the induction of Striga seed germination (Tsuchiya & McCourt, 2015). However, our findings showed a contradiction to this model because Striga seed exhibited germination independent of SL-ShHTL interactions. The internal reprogramming and regulation in hormonal balance by FL, especially reduction in ABA, and induction of gibberellin (GA) can recruit many physiological processes inside the seeds that could ultimately contribute to seed germination (Kusumoto et al., 2006; Machin and Bennett, 2020).

To further investigate the link between ABA, seed conditioning, and seed germination, we determined ABA content in unconditioned Striga seeds exposed for 24 h to water, FL, *rac*-GR24, or FL + *rac*-GR24. We also measured ABA in the corresponding seed exudates, assuming that the previously reported release of ABA by Striga seeds (Fujioka et al., 2019a) might be part of the conditioning/germination processes. By using Liquid Chromatography Tandem-Mass Spectrometry (LC-MS/MS), we detected a high level of ABA in H_2_O-treated Striga seeds and their exudates. Seeds treated with *rac*-GR24 showed a lower ABA content, but their exudates contained amounts comparable to those of H_2_O-treated seeds. As expected, the application of FL or FL + *rac*-GR24 caused a significant reduction of ABA content in both seeds and seed exudates (Figure 1b).

Previous result demonstrated that CYP707A protein could be the key determinant of ABA-dependent seed dormancy by catabolizing ABA (Brun et al., 2019); therefore we checked the transcript level of *CYP707A1* and *CYP707A2*, which were significantly induced in unconditioned seeds after 24h treatment with H_2_O, irrespective of chemical treatment (FL or *rac*-GR24). This result indicates that the water enhances ABA metabolism in Striga seeds after a 24 h short-term conditioning. In parallel, the 8 days of conditioning significantly enhanced the *CYP707A1* and *CYP707A2* expression compared to unconditioned seeds. Although 6 h *rac*-GR24 treatment fluctuates the *CYP707A1* expression in conditioned seeds, both expression level of *CYP707A1* and *CYP707A2* remained significantly higher than in unconditioned seeds, suggesting a higher ABA catabolic activity (Figure S2).

Taken together, Striga seeds release ABA during the early conditioning phase and that applying SLs at this stage does not have a pronounced effect on ABA release. Besides, FL suppresses ABA biosynthesis, shortening the conditioning phase, and enabling Striga seeds to break dormancy and germinate.

### 3.2 Sensitivity to ABA decreases during Striga seed conditioning

To further understand the role of ABA during the conditioning and germination processes, we conditioned the seeds in water for two or eight days and applied *rac*-GR24 (1.0 μM) alone or combined with various doses of ABA (1.5, 3, 6, 12, 25, 50, and 100 μM) for 24 h (Figure 1c). Inhibition of Striga seed germination by ABA was evident in 2 days conditioned seeds. The reduction in Striga seed germination was concentration-dependent that was reduced to around 66% upon the application of ABA at 100 μM concentration as compared to *rac*-GR24 control treatment (0 μM of ABA). However, 8 days conditioned Striga seeds showed a significantly lower sensitivity to ABA. Thus, the application of ABA at 100 μM concentration caused only a 22% reduction in seed germination (Figure 1c). These findings evidenced the role of ABA as a germination inhibitor, particularly in unconditioned Striga seeds; in contrast, in conditioned seeds, the ABA level was negligible before germination, while upon SL application, high concentrations of ABA were detected in the germinated seedlings. The amount of ABA increased drastically within twelve hours, indicating that Striga is able to synthesize ABA independently (Fujioka et al., 2019a). However, a decreasing trend of ABA levels in germinating Striga seeds (Toh et al., 2012) and P. ramosa seeds (Lechat, et al., 2012) were also observed that contradict to the recent finding (Fujioka et al., 2019a). Difference in parasitic spp., eco type, age of the parasitic seeds, and experimental conditions could be possible reasons behind these inconsistent results. The sensitivity of Striga seeds towards ABA decreases during the conditioning process, as seeds conditioned for 2 days were significantly more affected by ABA than the ones conditioned for 8 days (Figure 1c). This ABA-insensitive phenotype appeared in Striga to sustain its unique survival strategy. Indeed, the Striga ABA receptors stop regulating protein phosphatase 2C (ShPP2C1) due to a specific mutation in its amino acid sequence, interrupting ABA signalling in Striga. It helps Striga to enhance its transpiration rate for siphoning off water and therein dissolved nutrients from the hosts (Fujioka et al., 2019a; Fujioka et al., 2019b). Consistenly, we observed that *PP2C1* expression was upregulated in H_2_O-treated seeds after 24h (mock) compared to unconditioned seeds, whereas no difference was observed upon chemical treatments. Although conditioning and *rac*-GR24 treatment downregulate the *PP2C1* expression, no difference was observed in 6- and 24-h *rac*-GR24 treated seeds compared to conditioned seeds. Thus, our findings suggest that the *PP2C1* expression was neither responsive to *rac*-GR24 nor changing ABA content in Striga (Figure S2).

### 3.3 SL-induced seed germination triggers ABA biosynthesis and release

Next, we quantified the ABA content in unconditioned Striga seeds and in conditioned Striga seeds at 2-day intervals up to 8 days. Moreover, we treated 8 days conditioned Striga seeds with *rac*-GR24 (1.0 μM) for 6, 12, and 24 h and monitored ABA content both in seed tissues and seed exudates. Unconditioned Striga seeds showed high ABA content, which was significantly reduced when soaking in water from day 2 to onward. However, exposure of 8-days-conditioned seeds to *rac*-GR24 showed a gradual increase in ABA content in seeds and their exudates, showing the highest levels after 24 h of treatment (Figures 1d and 1e). To confirm the high release of ABA upon germination, we used Arabidopsis *pMKKK18::GUS* marker line as a sensor of this hormone (Mitula et al., 2015). As shown in (Figure 1f), exudates collected from seeds exposed to 24 h *rac*-GR24 treatment caused a strong induction of the GUS activity in the primary root of this Arabidopsis reporter line after 6 h of application. The GUS reporter induction at the primary root tip was stronger than that observed with ABA (50 μM) alone.

To support these findings, we further investigated the impact of SL ±FL on Striga emergence under greenhouse conditions in pots using the suicidal germination protocol (Jamil et al., 2022a; Jamil et al., 2022b). We initially tested the activity of FL and formulated SL analog (MP3 27EC) at 1.0 μM concentration under lab conditions (Figure 2); therefore, we decided to apply the same concentrations in the Striga emergence assays in the soil. For this purpose, we first induced germination of unconditioned Striga seeds in the infested pots with FL ± SLs for suicidal death for two weeks and then grew the rice (IAC-165) as a host crop to determine the Strgia emergence in each pot (Figure 2). Except the mock (untreated seeds), we applied the SL analog MP3 27EC (Jamil et al., 2022a) with or without FL as further controls. Untreated Striga-infested pots were included in the experiment as control, which showed the highest maximum Striga infestation (∼ 5 Striga per pot), accompanied by a severe reduction in the height of the host plant. Treatment with FL led to an apparent reduction in the Striga emergence; however, it also showed a significant negative impact on the host growth (Figure 2). Co-application of FL and MP3 led to similar results. Our results suggested that inhibiting ABA biosynthesis by FL in Striga seeds can trigger suicidal germination of Striga seeds and decrease their emergence at significant levels.

**FIGURE 2.**
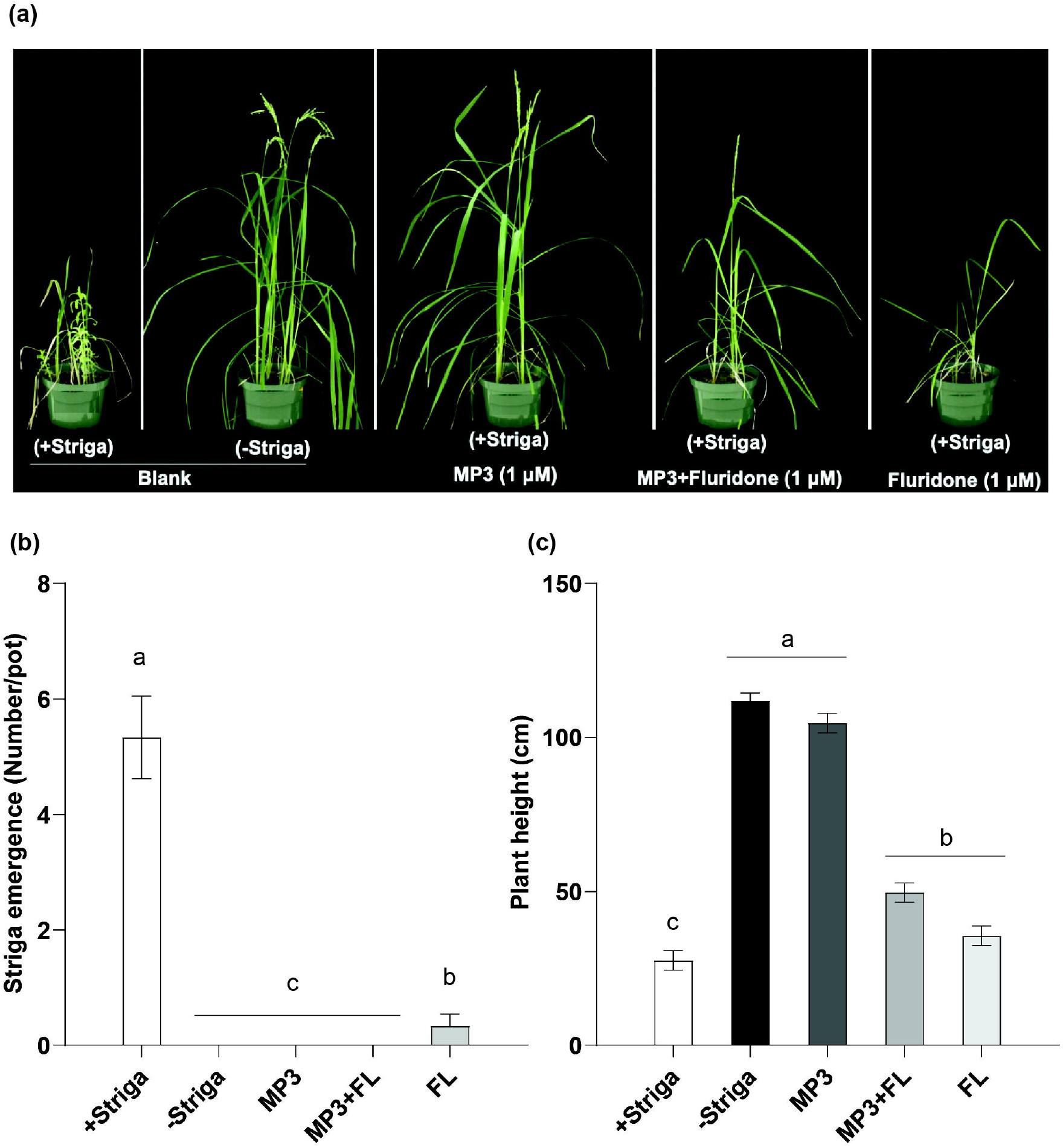
Striga emergence under greenhouse conditions in response to SL analogue MP3 and FL application. (a) View of Striga emergence in some representative pots under greenhouse conditions. Various treatments (MP3 ± FL at 1.0 μM each) at 500 ml per pot were applied twice on day 1 and day 3. (b) Striga emergence and (c) Rice plant height in response to various treatments. Data are means ± SE (n=6). The treatments with various letters denote significance (one-way ANOVA, Tukey’s post hoc test, P <0.05).

### 3.4 ABA released by germinated Striga seeds delays conditioning and germination of surrounding seeds

Taking into consideration its negative effect on seed conditioning and germination, we assumed that ABA released by germinated Striga seeds might negatively impact these processes in neighboring Striga seeds and have functions other than the previously reported “bewitching” of host plants (Fujioka et al., 2019a; Fujioka et al., 2019b). To test its effect on neighboring seeds, we placed discs with seeds at different stages of the conditioning phase (day 0 to 7) around a disc with seeds that have been germinated and exposed to *rac*-GR24 for 24 h. After one day, we treated the seeds undergoing conditioning phase with *rac*-GR24. One day later, we determined their germination rate (Figures 3a and 3b). We used discs soaked with ABA (100 μM) as a positive control and discs carrying 8-day-conditioned seeds treated with water as a negative control. Evaluation of the germination rates indicated that already germinated seeds inhibited the germination of seeds during the conditioning phase, due to the release of ABA. This effect was most pronounced on seeds that entered this experiment without conditioning (around 70%, day 0) and decreased with increasing conditioning time. The use of discs soaked with ABA caused complete inhibition of the germination of seeds without conditioning. Here again, the inhibitory effect decreased with increased conditioning time, as Striga seeds lost their sensitivity towards ABA during the conditioning phase (Figure 3b). These data suggest that ABA released by germinating Striga seeds negatively impacts the germination of the surrounding ones that are either unconditioned or within the conditioning stage, providing a competitive advantage in infesting host roots and maximizing the exploitation of its resources.

**FIGURE 3.**
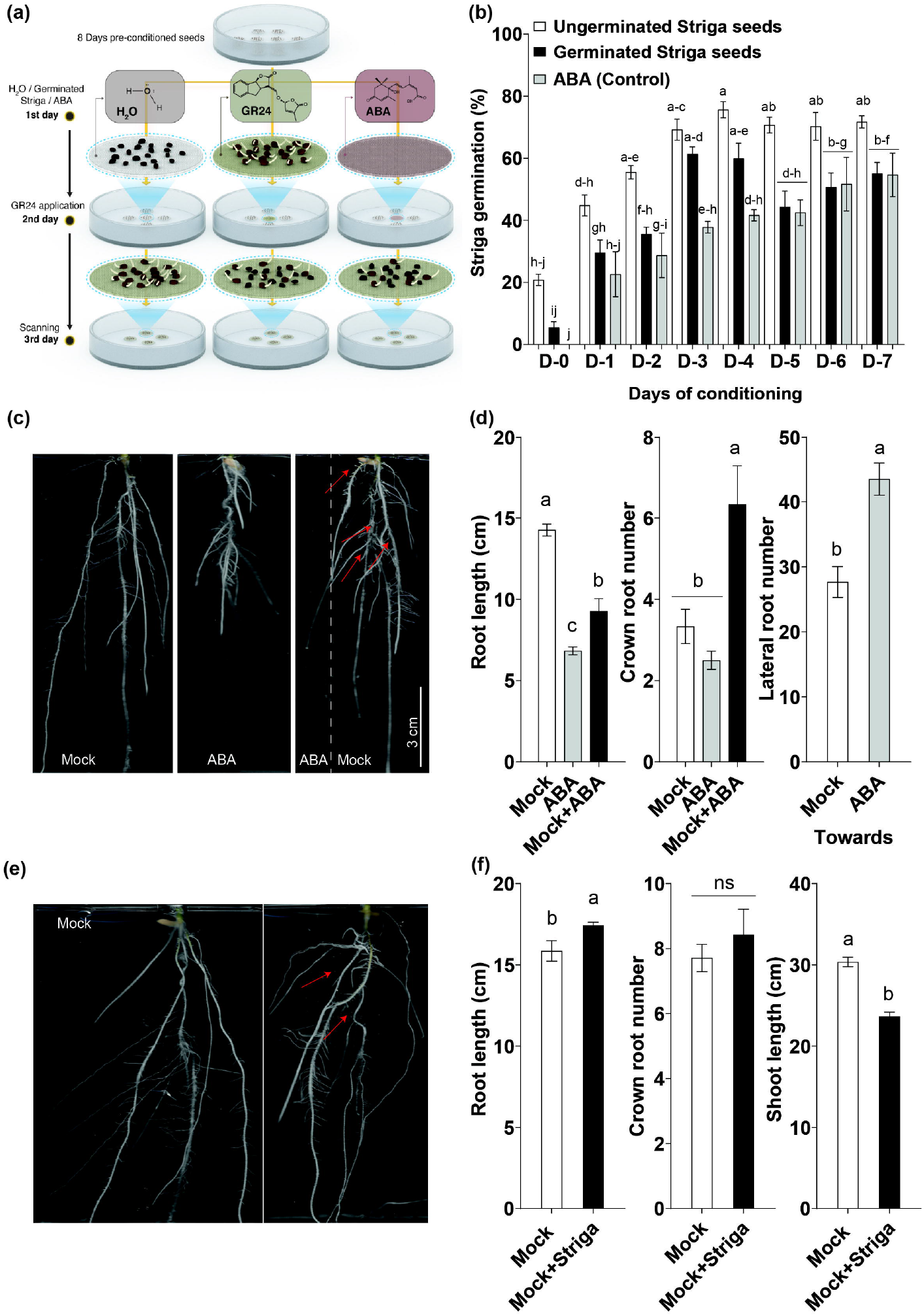
Impact of ABA released by germinated Striga seeds on the conditioning/germination of surrounding Striga seeds and the growth of host roots. (a-b) Effect of ABA released by germinated Striga seedlings on the germination of neighboring Striga seeds in the rhizosphere. Striga seed germination bioassays were performed with pre-conditioned (8 days) seeds pre-treated with water or 1.0 μM *rac*-GR24 for 24 h, placed on a disc, and used as an ABA inducer to modulate germination. The inducer discs were put each between four Striga discs containing seeds pre-conditioned for 0 to 7 days and incubated for another 24 h for ABA release. On the following day, we treated the surrounding four Striga discs with 1.0 μM *rac*-GR24 for 24 h, then, determined their germination rates. We used a disc soaked with 100 μM ABA as a positive control. Data are means ±SE (n=4-8). (c-d) Effect of ABA on host root growth. Rice seedlings (IAC-165) were treated in agar plates with Mock and/or ABA (5.0 μM) for one week to measure root length, crown root numbers, and lateral root numbers of roots growing towards the mock or ABA side. Each doc represents a biological replicate. Data are means ±SE (n=6). The treatments with various letters denote significance (one-way ANOVA, Tukey’s post hoc test, P <0.05). (e-f) Effect of pre-germinated Striga seeds on root/shoot growth of rice seedlings. (e) Root phenotype of un-infected and Striga-infected rice seedlings. (f) Root length, number of crown roots, and shoot length of rice seedlings (IAC-165) in the presence of germinated Striga seeds. Measurements were performed on the side exposed to Striga seeds. Each doc represents a biological replicate. Data represent means ±⍰SE (n⍰=⍰7). The treatments with various letters differ significantly according to one-way ANOVA and Tukey’s post hoc test (p <0.05). Error bars represent the standard error of the mean. ns: non-significant.

A further possible biological role of released ABA may be related to manipulating the host in a way increasing the probability of infestation. Indeed, it was reported that ABA released by germinating Striga seeds may lower the host’s immune system, making it more vulnerable to the parasitic attack (Fujioka et al., 2019a). Taking into consideration its known role as a determinant of root development (Miao et al., 2021), we hypothesized that ABA released by germinating Striga seeds might modulate the growth of the host’s roots to make them more accessible for infestation. To check this possibility, we first tested the effect of ABA. As expected, plants cultivated directly with 5.0 μM ABA showed root and shoot growth inhibition. However, we observed high numbers of crown and lateral roots in rice seedlings grown in agar plates supplemented on one side with the same concentration of ABA (Figures 3c and 3d), suggesting that ABA at low concentrations arising through diffusion from the ABA side promoted root growth (Figures 3c and 3d), which is in line with a previous report (Chen et al., 2006). Next, we determined the effect of germinated Striga seeds on host roots. For this purpose, we placed Striga seeds, 24 h after germinating them by *rac*-GR24, on one side of agar plates on which we grew rice seedlings. Interestingly, we observed a significant increase in root length and number of lateral roots, compared to the mock control (Figures 3e and 3f; Figure S3), indicating that ABA or unknown metabolites released by Striga seeds modulate host roots development. Although there might be unidentified compounds released by Striga during conditioning, we observed a reduction in shoot length of rice seedlings exposed to germinated Striga seeds, likely due to Striga-released ABA (Figures 3e and 3f). Taken together, we propose a model emphasizing the importance of ABA in modulating Striga seed conditioning and suicidal germination processes (Figure 4).

**FIGURE 4.**
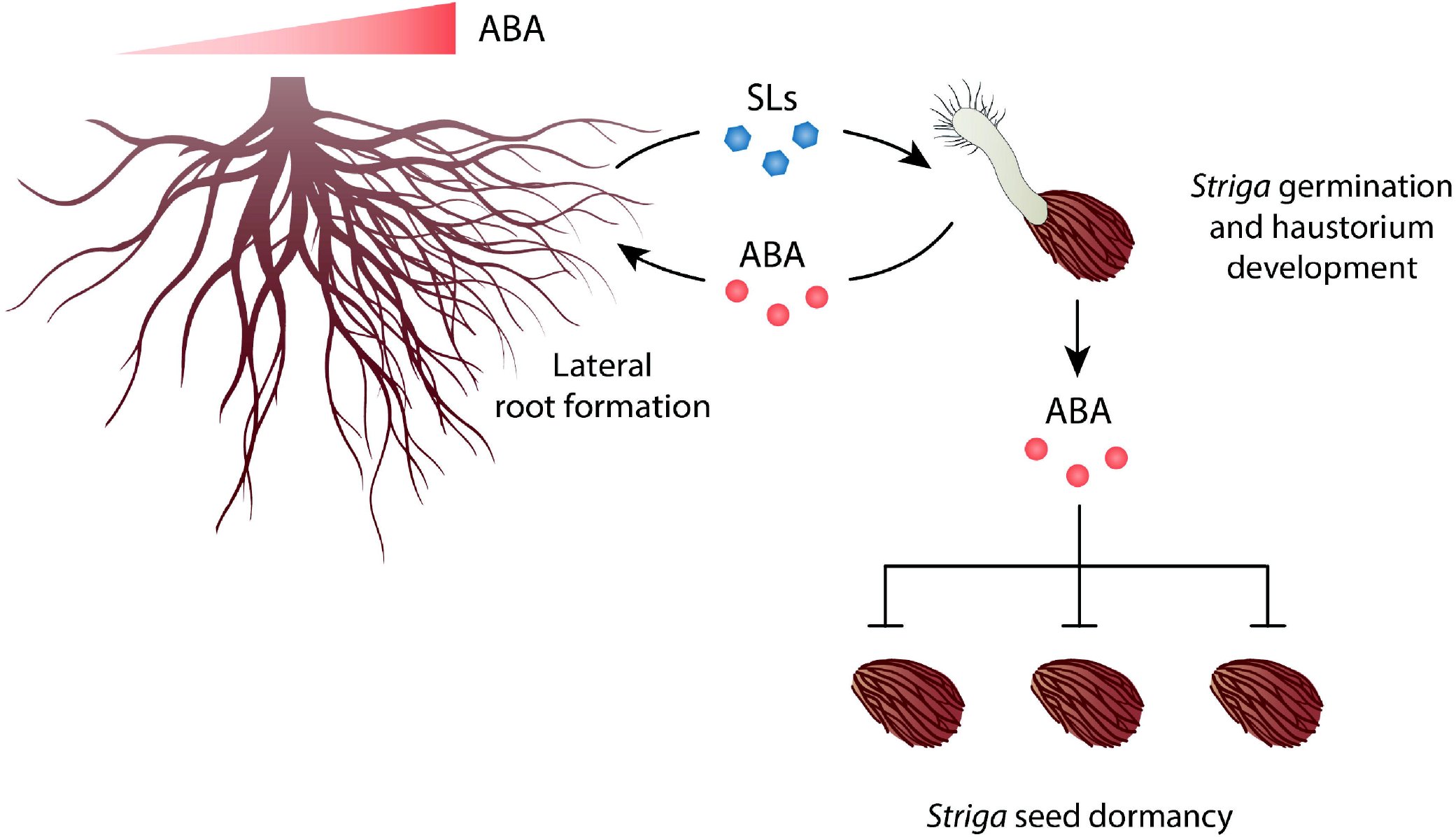
Proposed illustration and model showing the role of ABA in Striga germination and host infestation. Conditioned Striga seeds germinate upon perceiving host-released SLs. Germination initiates the development of a haustorium that enables host infestation and promotes ABA biosynthesis and release. ABA released into the rhizosphere negatively impacts the conditioning and germination of surrounding Striga seeds and promotes the growth of host roots towards the growing parasite, providing a competitive advantage and supporting host infestation.

In summary, we show that ABA is a major determinant of Striga seed conditioning and germination processes. ABA stored in Striga seeds is responsible for their prolonged dormancy that can be broken during conditioning in a moist and warm environment, making them able to perceive host-released germination stimulants. After a decrease in ABA content upon conditioning, induction of germination by SLs triggers ABA biosynthesis and release. We provide evidence that the increased ABA biosynthesis and release upon germination is coupled with decreased ABA sensitivity. Moreover, our data indicate a novel role of this plant hormone as a rhizospheric signal, in an SL-independent manner, serving in the Striga/Striga communication, i.e. between Striga seeds at different conditioning stages, providing a competitive advantage for already germinated seeds. In addition, we show that released ABA facilitates the infestation process by promoting the root growth of the host (Figures 4).

## Acknowledgments

This study was supported by the Bill & Melinda Gates Foundation (grant number OPP1136424) and baseline funding from King Abdullah University of Science and Technology given to S. A.-B. We are grateful to Late Prof. Abdel Gabar Babiker, The National Research Center, Sudan, for providing *Striga hermonthica* seeds. We are thankful to Dr Jonne Rodenburg, Africa Rice Center, Tanzania, for providing seeds of rice IAC-165. We are grateful to KAUST Research Publication Services and Research Communication (Carolyn Unck, Xavier Pita, Heno Hwang) for helping us to make figures for scientific illustration. We also grateful to Prof. Ikram Blilou, The Laboratory of Plant Cell and Developmental Biology, KAUST, for assistance in generating microscopic images.

## AUTHOR CONTRIBUTIONS

S.A.-B., M.J., Y.A., and J.Y.W. designed the experiments and coordinated the project. M.J. conducted Striga bioassays. Y.A. performed the GUS staining assays and gene rexpression analysis. J.Y.W., G.-T.E.C., and N.M.K. performed the metabolite analysis. J.Y.W., and G.- T.E.C. conducted rice root development experiment. M.J., J.Y.W., G.-T.E.C., L.B., conducted greenhouse experiment and H.N.J.K., and J.C.M. contributed intellectually. M.J., Y.A., J.Y.W., and S.A.-B. analyzed the data and prepared the figures. M.J. and J.Y.W. wrote first draft and all authors contributed extensively to the writing and editing and agreed on the final format of the manuscript.

## SUPPORTING INFORMATION

**FIGURE S1:** *In vitro* Striga germination in response to strigolactone analogs ± fluridone. The strigolactone analogs Methyl Phenlactonoate 3 and *rac*-GR24 were applied alone (at 1.0 μM) or along with fluridone (at 1.0 μM) on un-conditioned (for 72 h) and 10 days conditioned (for 24 h) Striga seeds and germination percentage was calculated. The bars represent means ± SE (n⍰=⍰5). The treatments with various letters differ significantly according to one-way ANOVA and Tukey’s post hoc test (p <0.05). Error bars represent the standard error of the mean.

**Figure S2:** The expression of key genes associated with ABA reception and catabolism in unconditioned and conditioned Striga seeds upon FL ±*rac*-GR24 application. Asterisks denote significance (one-way ANOVA, *p <0.05, **p <0.005, ***p <0.005, ****p <0.0005). Data are shown as ±STDEV (n=4).

**FIGURE S3:** Effect of pre-germinated Striga seeds ± ABA on root growth of rice seedlings. (a) Roots of rice seedlings co-cultured with Striga ± ABA (5⍰μM). (b) Root length (c) Crown roots and (d) Lateral roots of rice seedlings co-cultured with Striga ± ABA. Each data point represents one plant (n⍰=⍰8). Data represent means ± SE. The treatments with various letters differ significantly according to one-way ANOVA and Tukey’s post hoc test (p <0.05). Error bars represent the standard error of the mean.

**Table S1**. List of qRT-PCR primers used in this work.

