## Supplementary material for "Abscisic acid inhibits germination of Striga seeds and is released by them as a rhizospheric signal providing competitive advantage and supporting host infestation": Supporting Information.pptx

### Slide 1
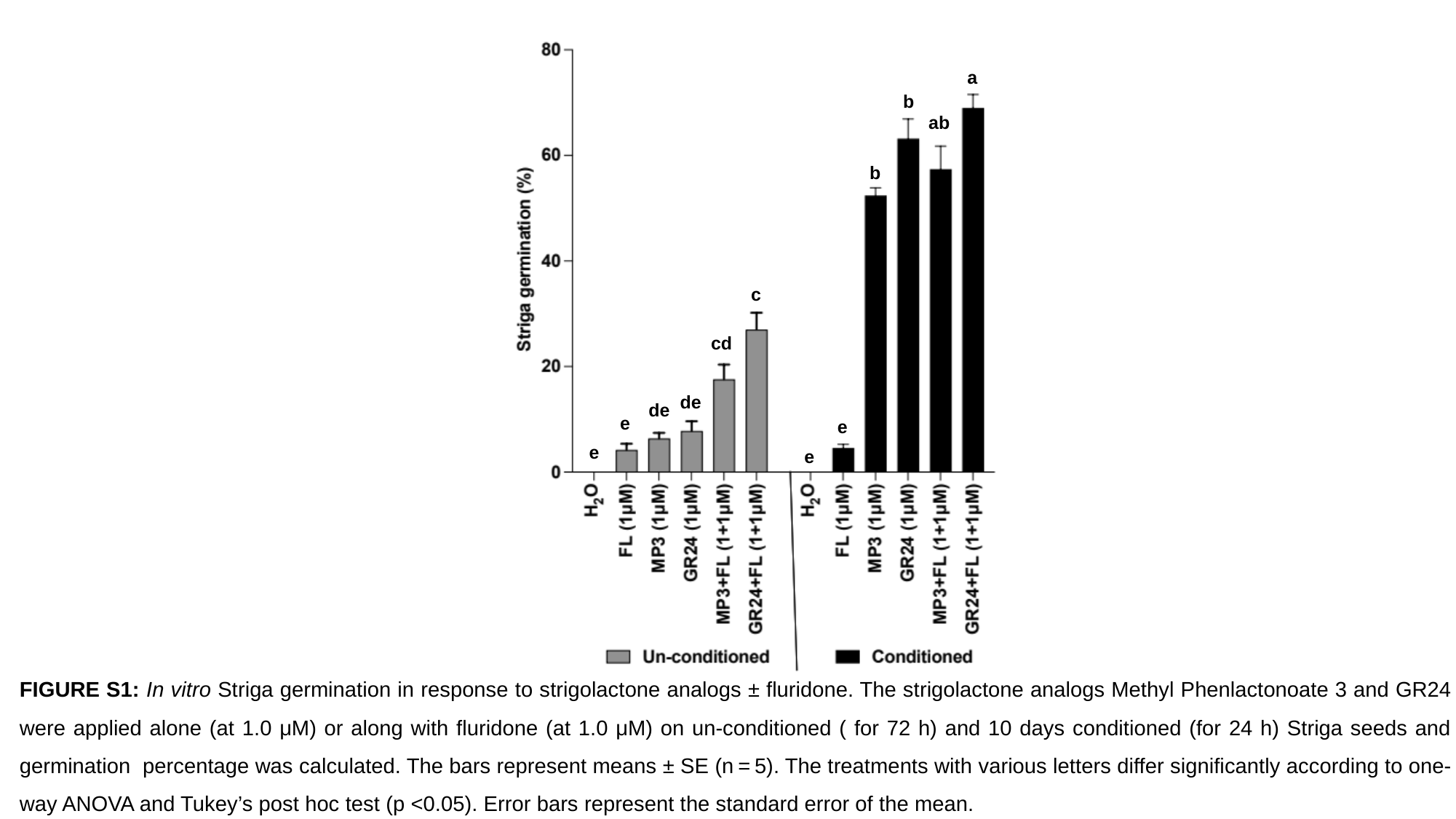

a
b
ab
b
c
cd
de
de
e
e
e
e
FIGURE S1: In vitro Striga germination in response to strigolactone analogs ± fluridone. The strigolactone analogs Methyl Phenlactonoate 3 and GR24 were applied alone (at 1.0 μM) or along with fluridone (at 1.0 μM) on un-conditioned ( for 72 h) and 10 days conditioned (for 24 h) Striga seeds and germination percentage was calculated. The bars represent means ± SE (n = 5). The treatments with various letters differ significantly according to one-way ANOVA and Tukey’s post hoc test (p <0.05). Error bars represent the standard error of the mean.

### Slide 2
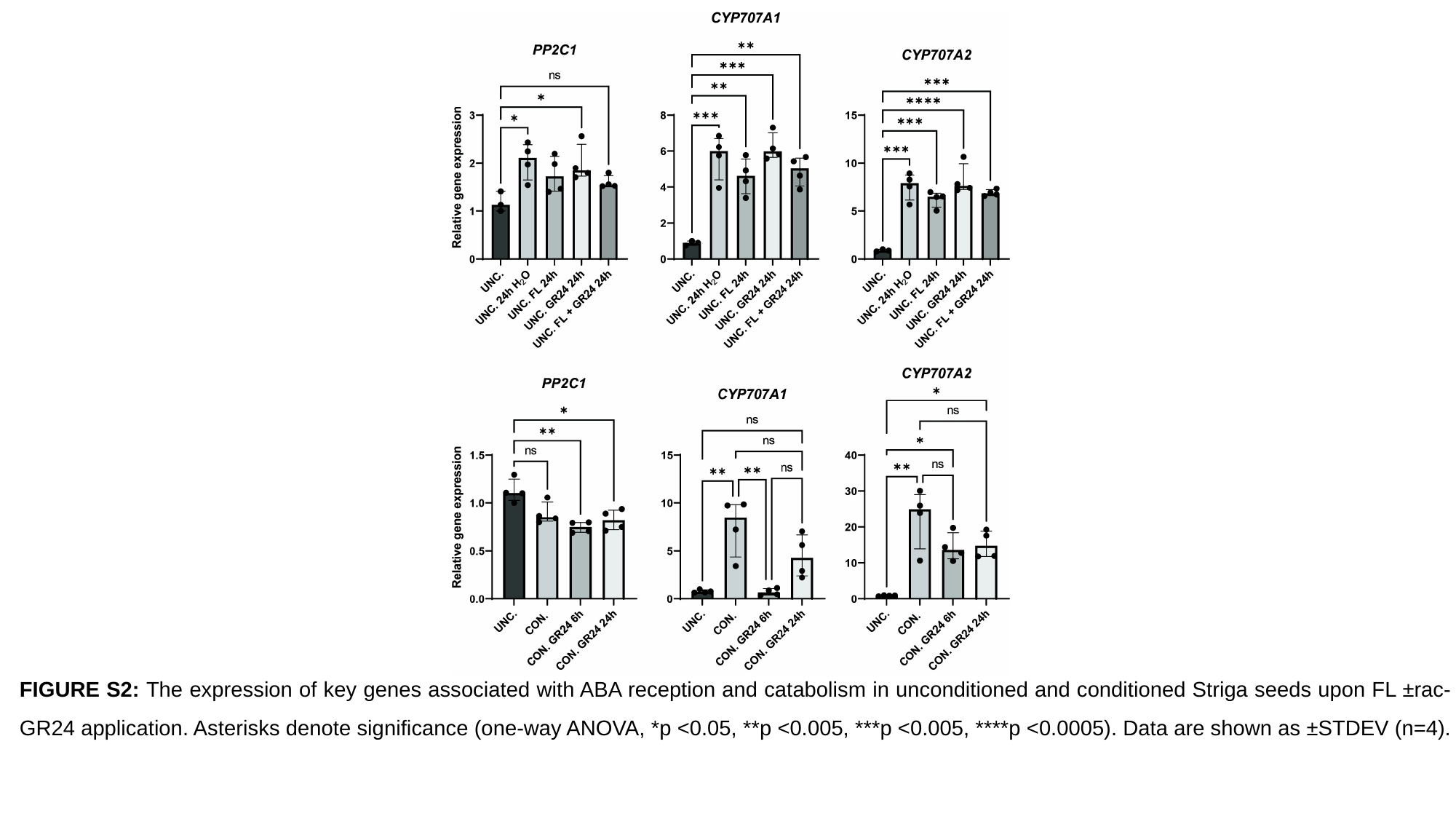

FIGURE S2: The expression of key genes associated with ABA reception and catabolism in unconditioned and conditioned Striga seeds upon FL ±rac-GR24 application. Asterisks denote significance (one-way ANOVA, *p <0.05, **p <0.005, ***p <0.005, ****p <0.0005). Data are shown as ±STDEV (n=4).

### Slide 3
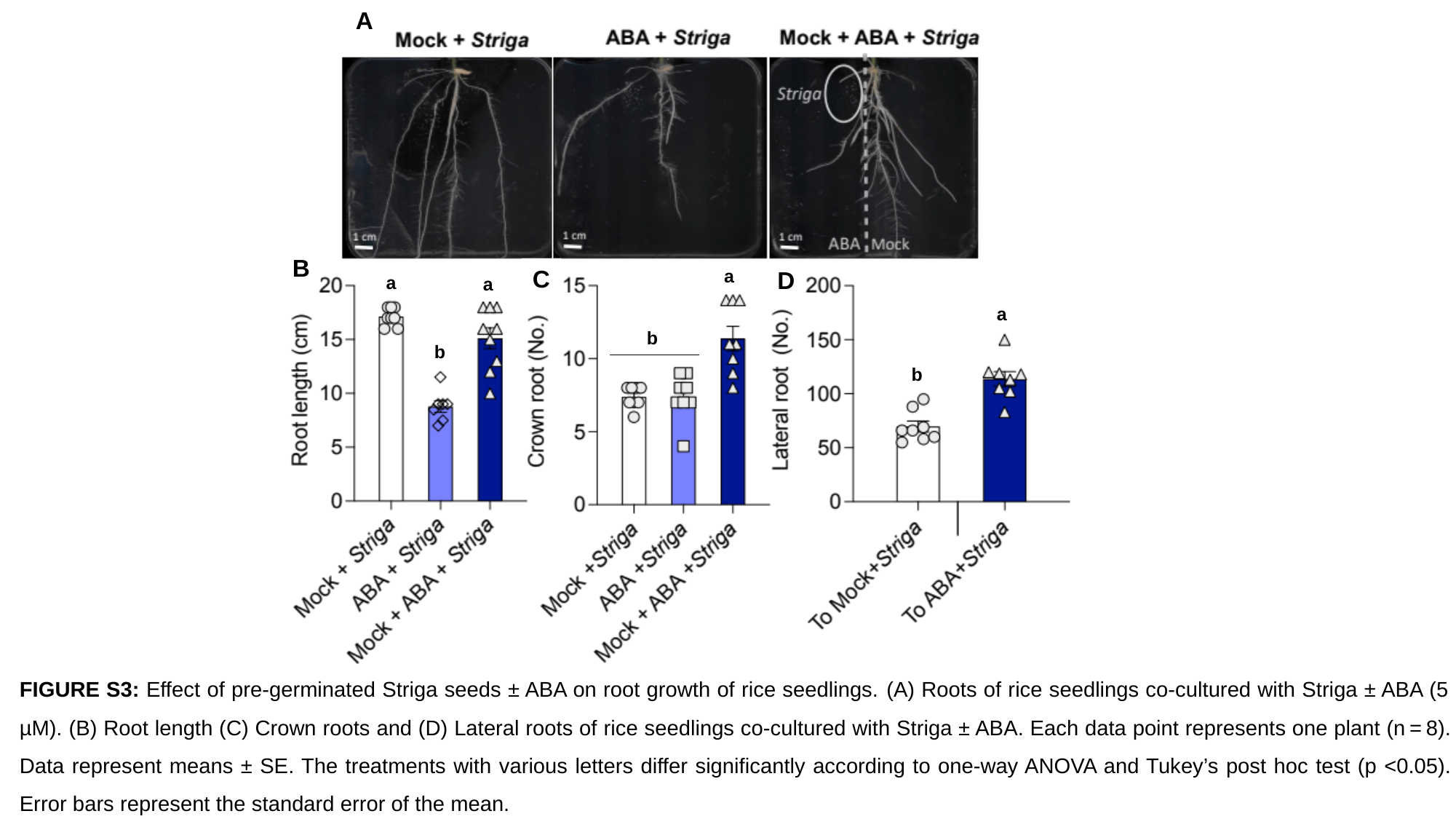

A
(b)
B
(c)
C
a
D
(d)
a
a
a
b
b
b
FIGURE S3: Effect of pre-germinated Striga seeds ± ABA on root growth of rice seedlings. (A) Roots of rice seedlings co-cultured with Striga ± ABA (5 µM). (B) Root length (C) Crown roots and (D) Lateral roots of rice seedlings co-cultured with Striga ± ABA. Each data point represents one plant (n = 8). Data represent means ± SE. The treatments with various letters differ significantly according to one-way ANOVA and Tukey’s post hoc test (p <0.05). Error bars represent the standard error of the mean.

### Slide 4
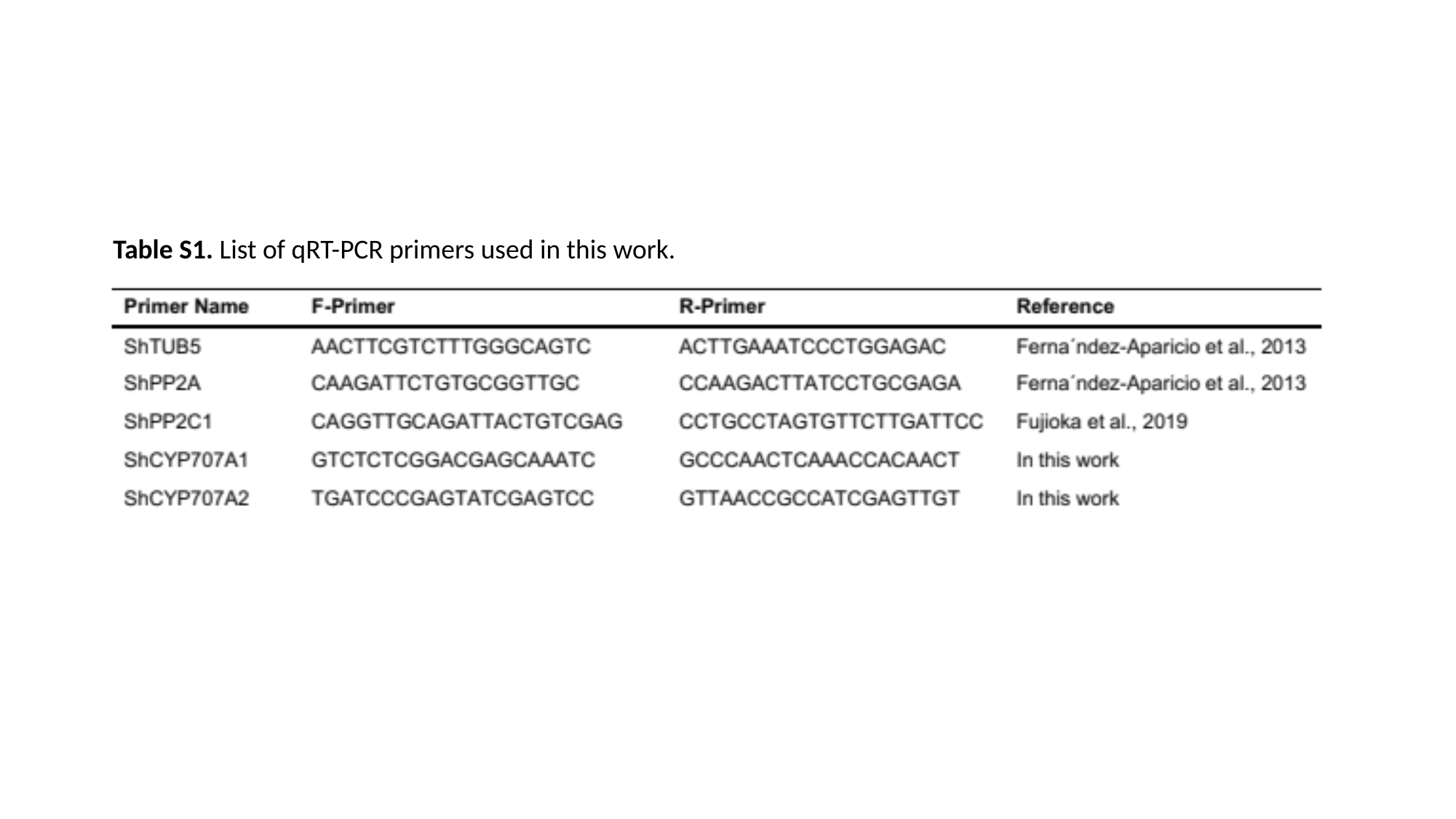

Table S1. List of qRT-PCR primers used in this work.
